# In vivo screen identifies LXR agonism potentiates sorafenib killing of hepatocellular carcinoma

**DOI:** 10.1101/668350

**Authors:** Morgan E. Preziosi, Adam M. Zahm, Alexandra M. Vázquez-Salgado, Daniel Ackerman, Terence P. Gade, Klaus H. Kaestner, Kirk J. Wangensteen

## Abstract

Existing drug therapies for hepatocellular carcinoma (HCC), including sorafenib, extend patient survival by only three months. We sought to identify novel druggable targets for use in combination with sorafenib to increase its efficacy. We implemented an *in vivo* genetic screening paradigm utilizing a library of 43 genes-of-interest expressed in the context of repopulation of the injured livers of Fumarylacetoacetate Hydrolase-deficient (*Fah*^−/−^) mice, which led to highly penetrant HCC. We then treated mice with vehicle or sorafenib to discover genetic determinants of sensitivity and resistance. Liver X Receptor alpha (LXRα) emerged as a potential target. To examine LXRα agonism in combination with sorafenib treatment, we added varying concentrations of sorafenib and LXRα agonist drugs to HCC cell lines. We performed transcriptomic analysis to elucidate the mechanisms of HCC death. *Fah*^−/−^ mice injected with the screening library developed HCC tumor clones containing *Myc* cDNA plus various other cDNAs. Treatment with sorafenib resulted in sorafenib-resistant HCCs that were significantly depleted in *Nr1h3* cDNA, encoding LXRα, suggesting that LXRα activation is incompatible with tumor growth in the presence of sorafenib treatment *in vivo*. The combination of sorafenib and LXR agonism led to enhanced cell death as compared to monotherapy in multiple HCC cell lines, due to reduced expression of cell cycle regulators and increased expression of genes associated with apoptosis. Combination therapy also enhanced cell death in a sorafenib-resistant primary human HCC cell line. Our novel *in vivo* screen led to the discovery that LXR agonist drugs potentiate the efficacy of sorafenib in treating HCC.

## INTRODUCTION

Hepatocellular carcinoma (HCC) is the third most common cause of cancer-related mortality worldwide and is increasing in incidence^1–3^. Intriguingly, a transcriptomic analysis of 17 different cancer types in humans revealed substantial overlap in all cancers with the exception of HCC^4^, which is consistent with the observation that treatments effective in other cancers have failed when applied to HCC^5^. HCCs are derived from hepatocytes^6^, and the unique characteristics of HCC may be related to the central role of hepatocytes in multiple metabolic processes including carbohydrate and fat storage and energy utilization, amino acid processing, bile salt synthesis recycling, protein detoxification, and drug metabolism, amongst other functions. Therefore, the development of new therapies will depend upon a better understanding and modeling of HCC.

Potentially curative interventions like liver transplantation are available to fewer than 30% of patients at the time of HCC diagnosis^7^. Sorafenib was the only first-line FDA-approved drug to treat HCC for over a decade^1, 8^. A multi-kinase inhibitor, sorafenib inhibits cell proliferation and angiogenesis through inactivation of various pathways including the ERK, VEGF, and PDGF cascades^9^. Resistance develops rapidly through numerous mechanisms leading to a median survival benefit of only 2-3 months. There are no good predictors of response to this treatment, likely due to inherent tumor heterogeneity^9, 10^. Furthermore, sorafenib has side effects including diarrhea, fatigue, and a syndrome of hand-foot skin reaction/rash, which often requires dose reduction^9^. Dual therapy with lower doses of sorafenib plus capecitabine^11^, doxorubicine^12^, and others^13, 14^, have not significantly improved survival and are not FDA-approved. Recently, additional multi-kinase inhibitors (e.g. cabozantinib) with mechanisms of actions similar to sorafenib, as well as anti-angiogenesis inhibitors and immunotherapy have been approved for treatment of HCC, but none of these have been shown to have improved survival over sorafenib^15–18^. Hence, there is still a need for broadly effective therapies.

Here, we report a conditional *in vivo* screen for drivers of tumor growth in the presence of sorafenib, designed to identify potential combination treatments for HCC. We validate a novel drug combination – sorafenib plus Liver X Receptor alpha (LXRα) agonist – with enhanced killing of HCC. LXRα agonists are currently in clinical trials and likely have acceptable side effect profiles for patients with advanced HCC.

## MATERIALS AND METHODS

### Animal tumor model

We previously described our method to perform genetic screening *in vivo* in the *Fah*^−/−^ mouse model of liver injury and repopulation^19^. These mice do not normally develop HCC unless oncogenes are provided^20^. *Fah*^−/−^ mice were maintained on 8 mg/liter nitisinone in the drinking water until the day of hydrodynamic tail vein injection (HTVI) with 10 μg of the plasmid library, consisting of equimolar amounts of the 44 different plasmids contained in the library (43 genes-of-interest and GFP). Each cDNA in the library has a unique 5-nucleotide barcode in the 3’ untranslated region to facilitate linkage of cDNAs to tumors via high-throughput sequencing. After HTVI and removal of nitisinone, mice were monitored for weight changes and overall body condition score. Sorafenib (30 mg/kg) or a DMSO control solution was administered by gavage daily beginning 6 weeks post-HTVI and continuing to 15 weeks. Magnetic resonance imaging (MRI) was performed at the Penn Small Animal Imaging Facility immediately after euthanasia. All procedures were approved by our institutional animal care and usage committee.

### Tumor sequencing

Whole livers from mice receiving the plasmid library were fixed overnight in 4% paraformaldehyde, mounted in paraffin, serially sectioned, and stained with hematoxylin and eosin by the University of Pennsylvania Molecular Pathology and Imaging Core. Hematoxylin and eosin-stained sections were submitted to the University of Pennsylvania School of Veterinary Medicine Comparative Pathology Core for histological grading and calling of HCC lesions based upon guidelines^21^, and the pathologist was blinded to study conditions. The HCC tissue from unique lesions was collected from adjacent serial sections using a fine needle and a stereo microscope. This tissue was used directly for two rounds of PCR using primers flanking the area of the 5-nucleotide barcode as described previously^19^. Sequencing libraries from each tumor received a unique Illumina index to enable multiplexing, and sequencing was conducted on an Illumina MiSeq by the University of Pennsylvania Next Generation Sequencing Core. A detailed protocol is available upon request.

### Cell culture

Hep3B, Hepa1-6, HepG2, and AML12 cell lines were purchased from ATCC. Huh7 were purchased from JCRB Cell Bank through Sekisui XenoTech company. Hep3B, Hepa1-6, HepG2, and Huh7 were cultured in DMEM with 10% fetal bovine serum and 5% Penicillin/Streptomycin. AML12 was cultured in DMEM:F12 supplemented with 10% FBS, 10 μg/ml insulin, 5.5 μg/ml transferrin, 5 ng/ml selenium, and 40 ng/ml dexamethasone. The PGM898 cell line is a patient-derived cell line that was generated by xenografting immunodeficient mice with percutaneous core biopsy HCC tissue, then culturing dispersed cells from the tumor (University of Pennsylvania Institutional Review Board-approved observational clinical trial #825782; Ackerman, D., et al., *manuscript in preparation*). The patient was a male in his 50s with HCV cirrhosis and BCLC stage B HCC. Pathology of biopsies taken at the time of sample procurement demonstrated moderately differentiated HCC. After establishment of the cell line, cells were propagated on Matrigel-coated plates in DMEM:F12 with 10% FBS, 5% P/S, Glutamax, Hepes, and 10 μM ROCK inhibitor.

### Drug treatments and crystal violet staining

Sorafenib, GW3965, and T0901317 were prepared in DMSO. For drug treatments, 9 × 10^4^ cells were seeded in wells of a 24 well plate (or 10^5^ AML12 cells in 6-well plates), and the following day the drugs or DMSO were added in fresh media. Forty-eight or 72 hours after treatment, cells were washed with PBS, fixed with 4% PFA, and stained with 0.05% crystal violet for 20 minutes. Once plates were washed, dried and imaged, methanol was added and absorbance was measured at 540nm.

### RNA sequencing

Huh7 and Hep3B were seeded at 6 × 10^5^ cells in 6-well plates. Wells were treated with DMSO, 2 μM sorafenib, 5 μM GW3965, or 2 μM sorafenib plus 5 μM GW3965 (combination group). Twenty-four hours later, RNA was extracted (Qiagen RNeasy mini kit) and purity was confirmed using an Agilent Bioanalyzer. Following enrichment of mRNA using magnetic bead isolation of poly(A) sequences (NEB #E7490), we prepared sequencing libraries using a commercial kit (NEB #E7770S). Sequencing was performed on a HiSeq 4000. Differential expression analysis was performed using edgeR (R software). To create a heatmap, we calculated the Log2 fold change (Log_2_FC) of each sample normalized to the average values in the DMSO treatment group, then subtracted the median Log_2_FC value of the DMSO group. This DMSO-centered Log_2_FC was plotted using ComplexHeatmap (R software). Gene ontology (GO) pathway analysis was performed using package STRINGdb, and visualized with ggplot2 and stringr (R software). RNA-seq data was submitted to the Gene Expression Omnibus (GEO, accession # pending).

### RT-PCR analysis

PGM898 cells were treated for 24 hours and RNA was extracted as described above. Reverse transcription was performed with the High-Capacity cDNA Reverse Transcription Kit (Thermo-Fisher). qPCR was performed using Sybr green and the following primers: GAPDH Forward: GTCTCCTCTGACTTCAACAGCG, GAPDH Reverse: ACCACCCTGTTGCTGTAGCCAA, B2M Forward: CCACTGAAAAAGATGAGTATGCCT, B2M Reverse: CCAATCCAAATGCGGCATCTTCA, GADD45B Forward: GADD45B Reverse, FASN Forward: CTTCCGAGATTCCATCCTACGC, FASN Reverse: TGGCAGTCAGGCTCACAAACG, CDK2 Forward: ACCGAGCTCCTGAAATCCTC, CDK2 Reverse: CCACTTGGGGAAACTTGGCT, PCNA Forward: CCATCCTCAAGAAGGTGTTGG, PCNA Reverse: GTGTCCCATATCCGCAATTTTAT, CCNE1 Forward: CATCATGCCGAGGGAGCG, CCNE1 Reverse: AGGCTTGCACGTTGAGTTTG. For analysis, threshold cycle values were normalized to the average threshold cycle for GAPDH and B2M.

### Statistical analysis

All scatter plots and bar graphs independent of RNA sequencing data were created using GraphPad. Statistical analyses were performed using a two-way ANOVA with Tukey’s test for multiple comparisons, a one-way ANOVA with multiple comparisons, or a Student’s *t*-test, as distinguished in the figure legends.

## RESULTS

### Generation of an HCC model using a plasmid library

We previously engineered a library of 43 cDNAs important in liver growth and function, which are expressed from plasmids also encoding the FAH enzyme. This plasmid pool was hydrodynamically injected into the tail vein of *Fah*^−/−^ mice to identify drivers of liver repopulation^19^. We previously identified MYC as a potent driver of liver repopulation, and TNFR1 as a strong negative regulator of liver repopulation. Liver tumors develop rapidly in this model, as evidenced by the fact that plasmid library-injected *Fah*^−/−^ mice developed neoplasms as early as six weeks post-injection (Figure 1A, B). The tumors displayed significant heterogeneity and were confirmed to be HCC by a veterinary pathologist. We determined the putative HCC driver genes present in each tumor by performing high-throughput sequencing of the linked DNA barcodes present in isolated tumor nodules (Figure 1C, D). We found an median of 4 distinct cDNAs per HCC tumor (range 1-11). Moreover, when we sequenced DNA from multiple parts of the same tumor, we found the same cDNA composition, indicating that tumors are clonal (Figure 1D, black box). Remarkably, all HCCs contained the *Myc* cDNA, indicating that MYC was the most potent driver of tumorigenesis among all the genes tested.

**Figure 1:**
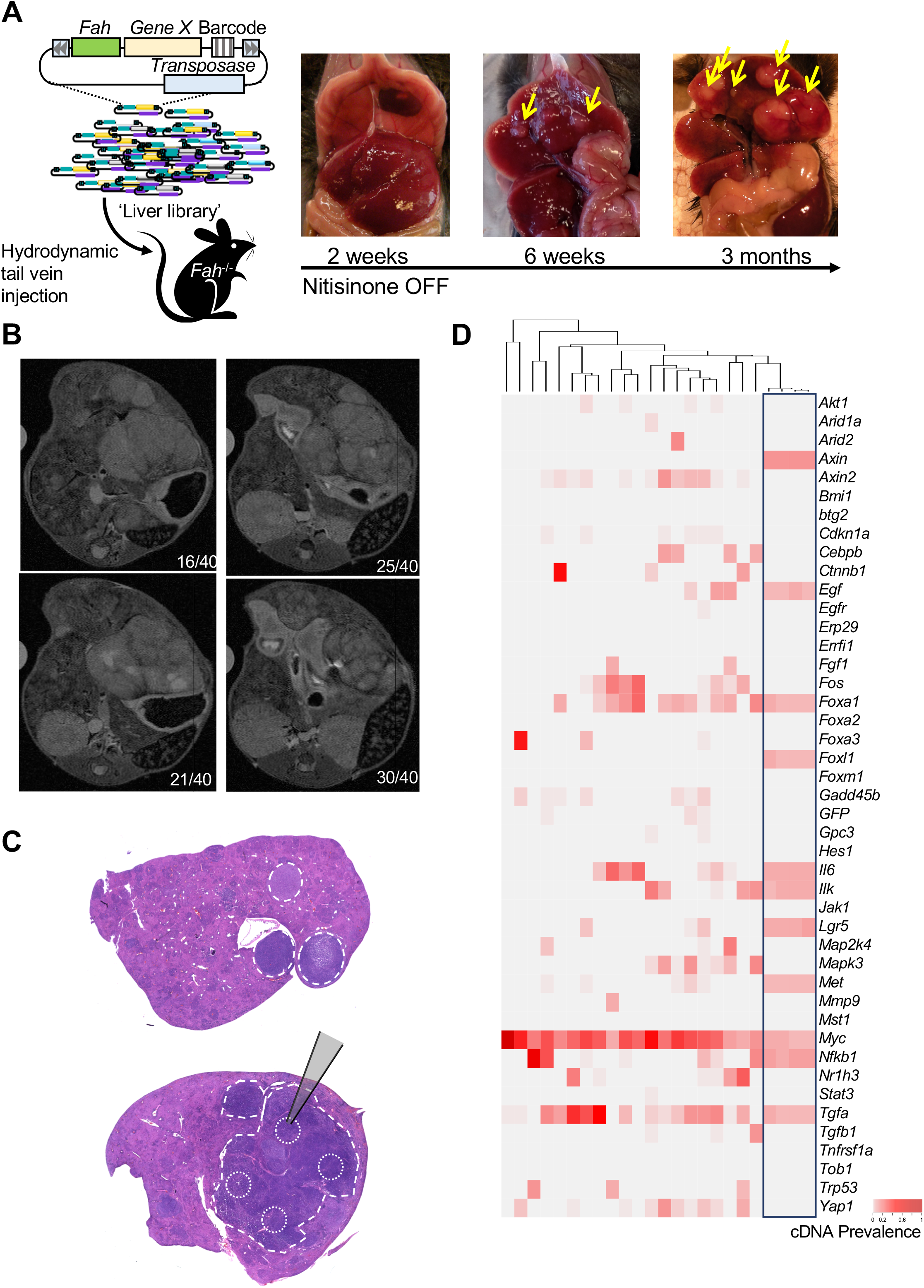
Generation of HCCs using a cDNA expression screen. (A) We injected a library of 43 genes-of-interest plus GFP, all linked to Fah expression, into *Fah*^−/−^ mice. We observed tumor growth by 6 weeks post-injection, and massive tumors by 3 months (representative image shown, N=3 mice at 6 weeks and N=4 at 3 months). (B) Representative MRI images at the time of euthanasia, showing multiple tumor masses in the liver. (C) Representative Hematoxylin and Eosin staining with tumor borders outlined. (C and D) Microdissected tumor DNA was amplified and sequenced to determine which barcodes, corresponding to specific cDNAs, are linked to the tumors (N=21 discrete tumor nodules from N=4 mice, range 3-10 tumors dissected per mouse). Each tumor had a median of 4 unique cDNAs from the library (range 1-11 with cut-off prevalence > 0.05). The four samples on the right side of the heatmap, outlined in black, were taken from separate parts of a single large tumor. The nearly identical pattern and close clustering on the cladogram indicate that the tumors are clonal.

### Sorafenib-resistant tumors cannot grow in the presence of *Nr1H3* (encodes LXR)

We hypothesized that our screening approach could be used to discover genes that confer sensitivity or resistance to sorafenib. Therefore, we injected *Fah*^−/−^ mice with the plasmid library, and allowed the mice to repopulate their livers for six weeks, the time-point at which early tumors are established. We then began daily treatment with either 30 mg/kg sorafenib or vehicle control for two months (Figure 2A). Sorafenib treatment significantly reduced the liver weight to body weight ratio and tumor burden compared to vehicle (Figure 2B, C). Intriguingly, sorafenib-resistant tumors had a higher mitotic index than vehicle tumors (Figure 2C).

**Figure 2:**
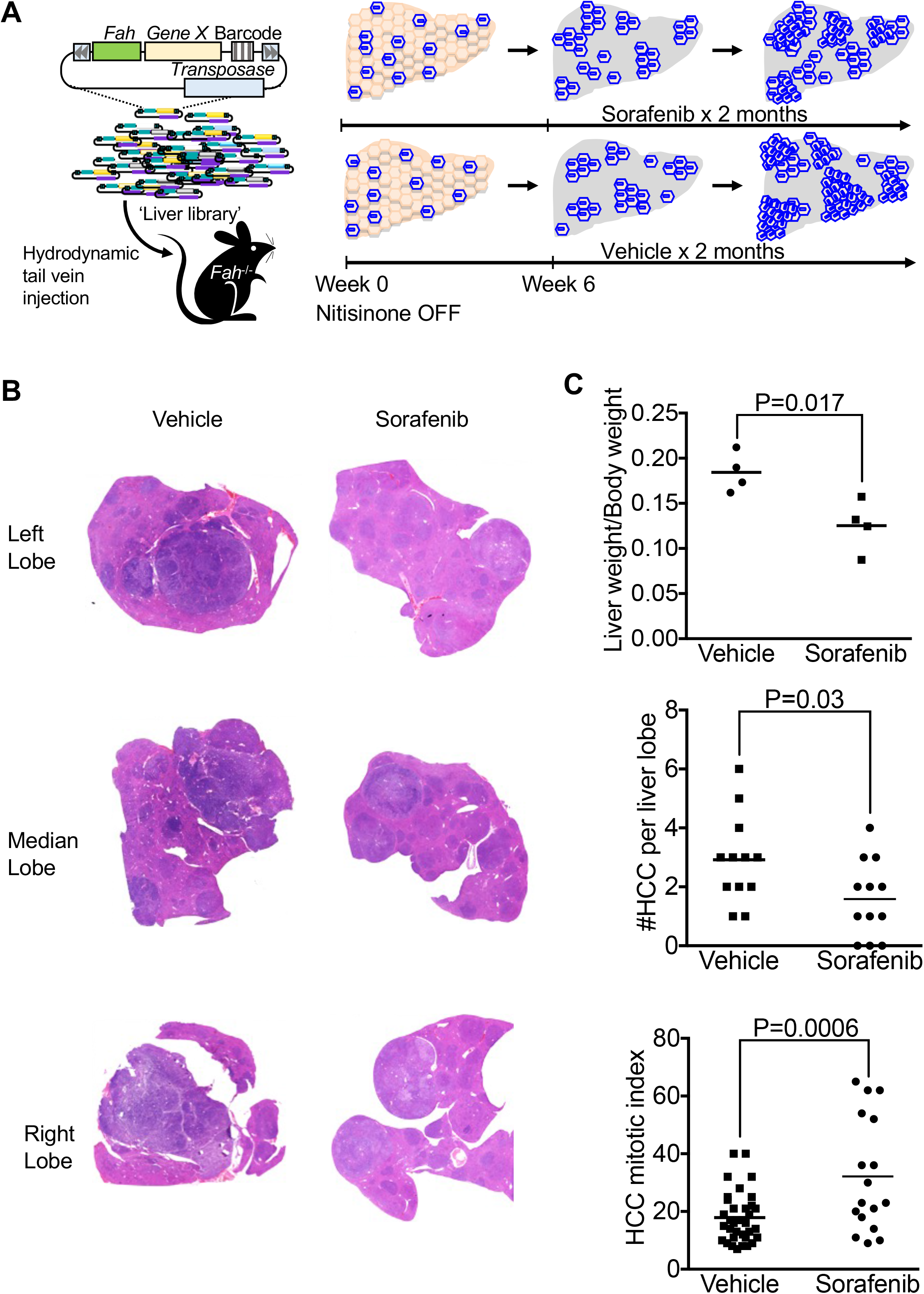
Sorafenib treatment reduces tumor burden. (A) Schematic representation of tumor generation and sorafenib treatment. (B) Representative Hematoxylin and Eosin staining of sorafenib- and vehicle-treated tumors. (C) Liver weight to body weight ratio, number of HCC tumors per liver lobe, and mitotic index of sorafenib- and vehicle-treated tumors (N=4 mice in each group). Statistics performed using Student’s *t*-test.

To identify genes correlated with sorafenib sensitivity, we analyzed which cDNA(s) were present or absent in HCCs that developed in the presence of sorafenib. Strikingly, we detected the *Myc* oncogene in all vehicle- and sorafenib-treated tumors assessed. However, the *Nr1h3* transgene, which encodes LXRα, was present in a number of vehicle-treated tumors as expected (Figure 1D, 3A). Intriguingly, it was completely absent from sorafenib-resistant tumors (Figure 3A, B). These results indicate that high LXR activity is incompatible with MYC-driven tumor growth in the presence of sorafenib.

**Figure 3:**
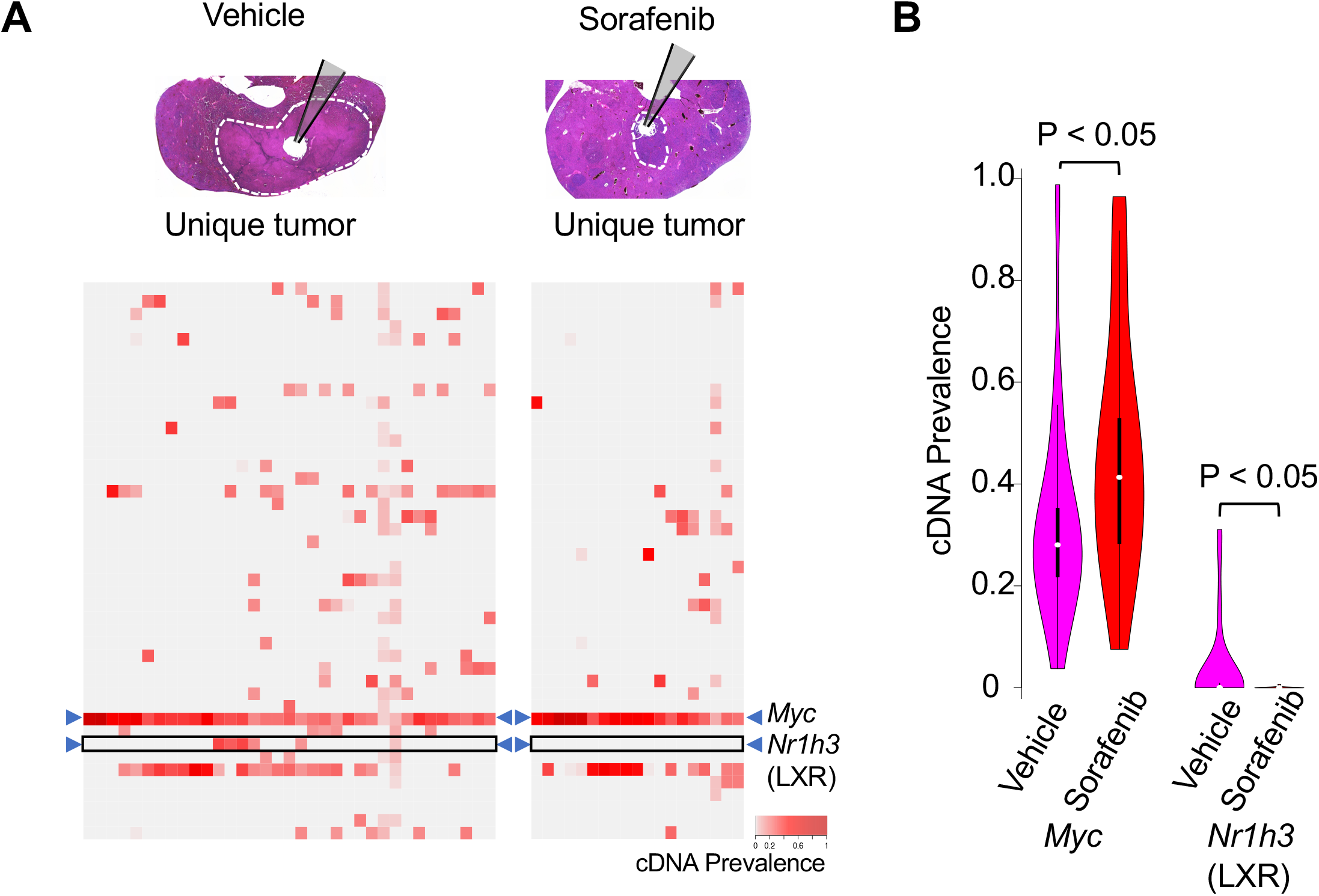
*NR1H3* (encoding Liver X Receptor, LXR) is incompatible with tumor growth in the presence of sorafenib. (A) HCC tumors were microdissected and sequenced to determine which cDNAs are linked them (N=35 vehicle-treated and N=19 sorafenib-treated HCCs). All sorafenib-treated and vehicle-treated tumors contained *Myc* cDNA. A subset of vehicle-treated tumors contained *Nr1h3* cDNA (which encodes LXR), while the cDNA was absent from the sorafenib-treated tumors (marked by a black box). (B) Violin plots showing sorafenib-treated tumors had high levels of linked *Myc* cDNA, and had complete absence of *Nr1h3*. Statistics performed using Student’s *t*-test.

### LXR agonists improve sorafenib efficacy against HCC *in vitro*

Based on our *in vivo* results, we asked whether activating LXR within cells using agonist drugs would improve response to sorafenib. We treated three human hepatoma cell lines (Hep3B, HepG2, and Huh7) and one murine hepatoma cell line (Hepa1-6) with varying doses of sorafenib and the small-molecule LXR agonist GW3965. We quantified cell viability after 48 hours by crystal violet staining (Figure 4). In each cell line, there was a 30-50% reduction in cell viability with the combination treatment at a minimal dose of 5 μM GW3965 and 2 μM sorafenib compared to sorafenib-only treatment, while treatment with 5 μM GW3965 alone did not affect cell viability (Figure 4B, D, E, G). At these doses, neither GW3965 nor sorafenib treatment alone impacted the murine hepatocyte-derived cell line AML12, whereas combination treatment led to only a 20% reduction from baseline (Supp. Figure 1A, B), suggesting treatment preferentially affects HCC cells. We also confirmed our findings in HepG2, Hep3B, and Huh7 cells using a second LXR agonist, T0901317, confirming that activation of LXR enhances sorafenib activity (Supp. Figure 1C, D). Taken together, these results suggest that LXR activation improves the efficiency of sorafenib in multiple HCC cell lines.

**Figure 4:**
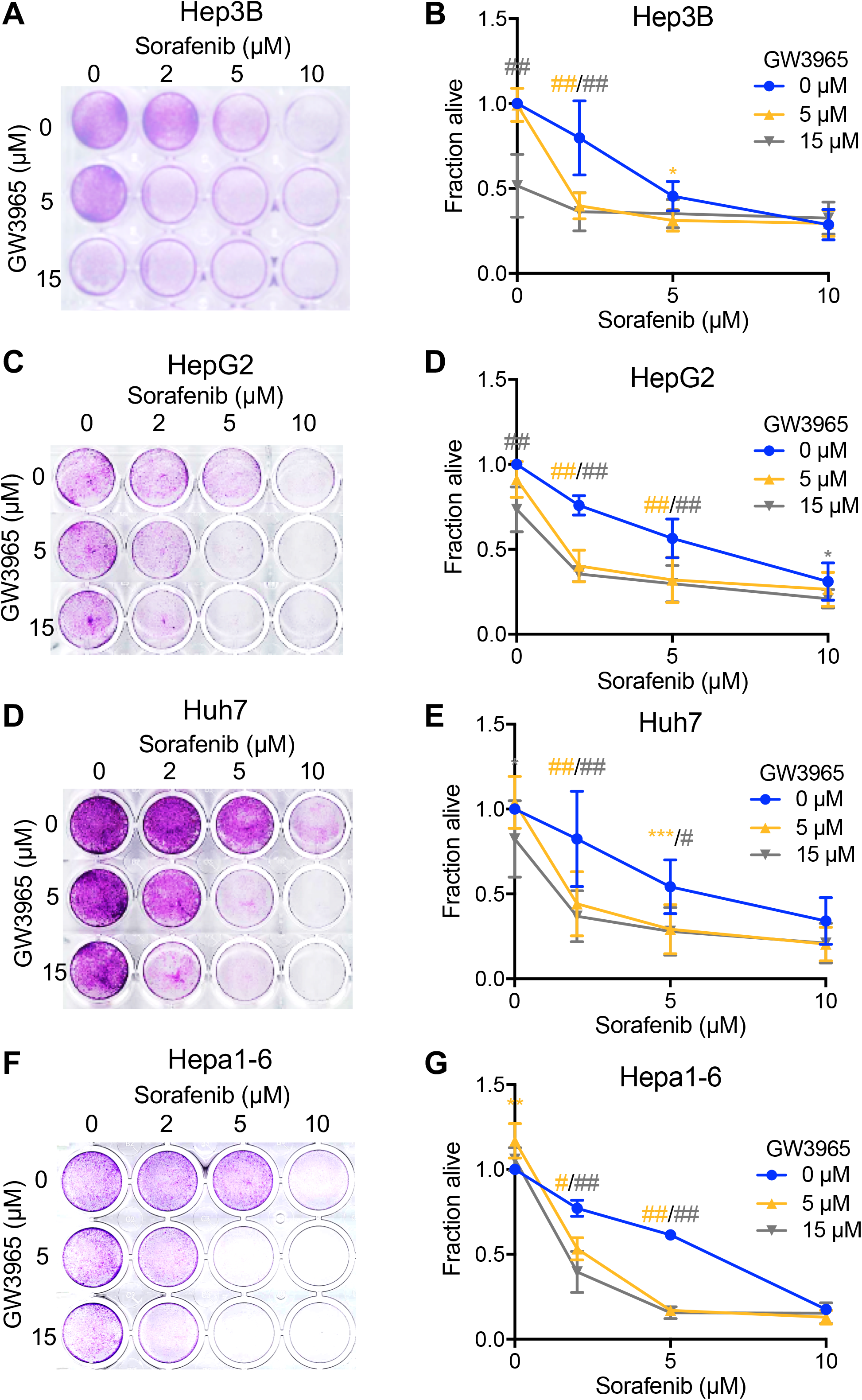
LXR agonist GW3965 and sorafenib combination treatment effectively targets HCC *in vitro*. Representative images and quantification of crystal violet staining of Hep3B (A, B, n=8), HepG2 (C, D, n=11), Huh7 (D, E, n=11), and Hepa1-6 (F, G, n=3) cells treated with varying concentrations of DMSO control, GW3965, sorafenib, or GW3965 and sorafenib for 48 hours. Quantification was performed by addition of methanol to plates and measuring absorbance (right). Statistical analysis performed with two-way ANOVA with Tukey’s multiple comparison. P values are as follows: *P<0.05, **P<0.01, ***P<0.005, #P<0.001, #P<0.0001.

### GW3965 and sorafenib work synergistically to target numerous pathways *in vitro*

We queried the mechanism by which the combination treatment targets HCC, and performed RNA sequencing in Hep3B and Huh7 cells treated with drugs for 24 hours. Principle component analysis identified clear separations between cell lines and treatment groups (Figure 5A). Next, we performed differential expression analysis comparing treatment to DMSO control groups, using a cut-off fold change of >2 (Log_2_FC >1 or <−1, Figure 5B). We found many genes to be altered by treatment in both cell lines congruently: 19 genes were differentially expressed in response to GW3965, 190 to sorafenib, and 915 to the combination treatment, overall highlighting a remarkable synergistic effect as visualized in Figure 5B.

**Figure 5:**
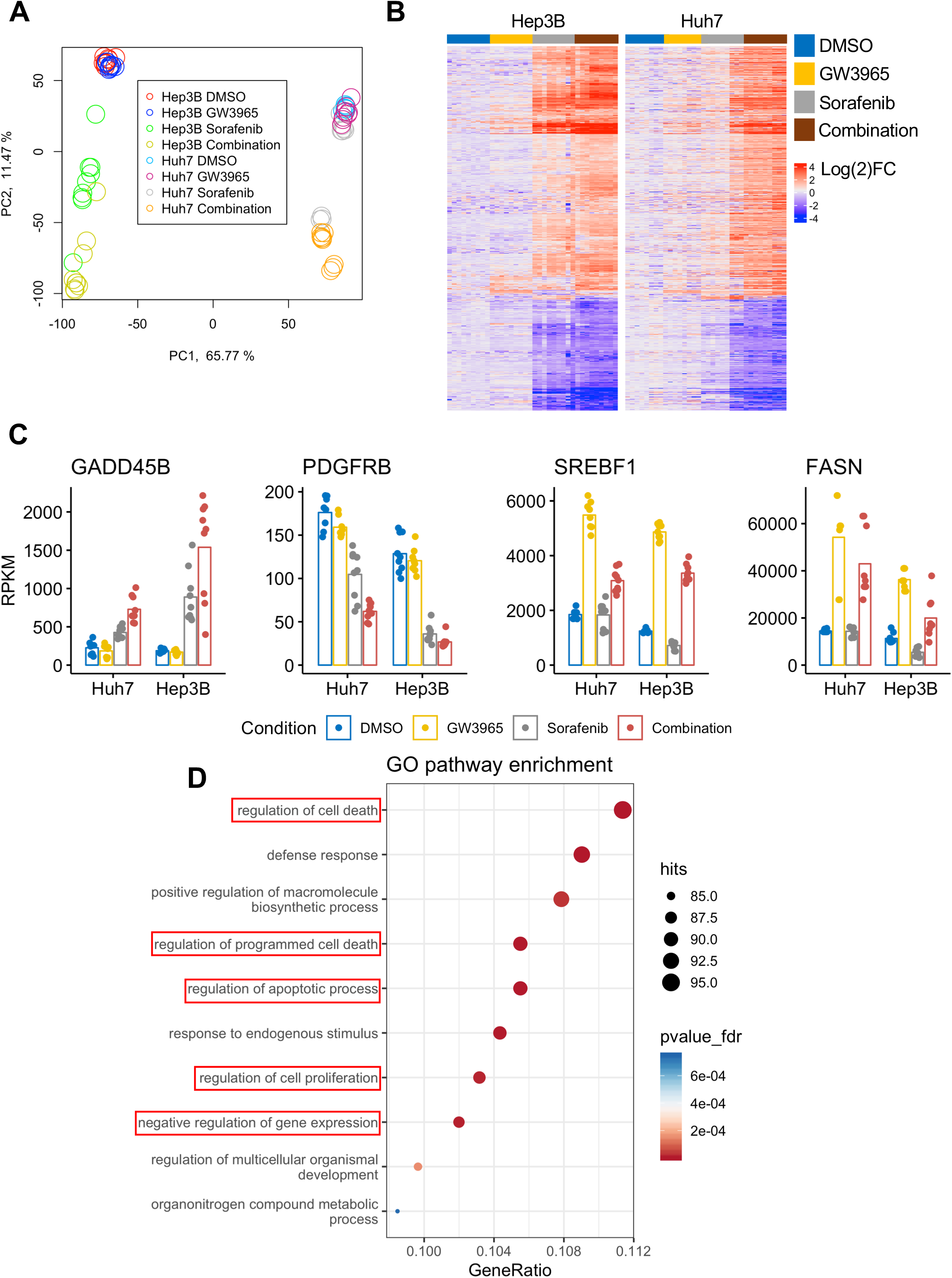
Transcriptomic analysis of drug treatments on HCC *in vitro*. Hep3B and Huh7 cells were treated with DMSO, 2 μM GW3965, 5 μM sorafenib, or GW3965 and sorafenib for 24 hours and processed for RNA sequencing (n=8-9). (A) Principle component analysis demonstrating Hep3B and Huh7 cluster separately on one axis, while the treatment groups cluster separately on the other axis. (B) Heatmap of differentially expressed genes. (C) Reads Per Kilobase of transcript, per Million mapped reads (RPKM) of Growth Arrest And DNA Damage-Inducible Protein GADD45 Beta (*GADD45B*), Platelet Derived Growth Factor Receptor Beta (*PDGFRB*), Fatty Acid Synthase (*FASN*), and Sterol Regulatory Element Binding Transcription Factor 1 (*SREBF1*). (D) Top 10 of 742 significantly modulated gene ontology (GO) pathways. Red boxes highlight pathways associated with the cell cycle or apoptosis.

Next, we confirmed the specificity of sorafenib and GW3965 treatment by assessing gene expression levels of known markers of drug activity. Upregulation of *GADD45B* and downregulation of *PDGFRB* mRNA levels are indicative of sorafenib activity^22, 23^, and we observed these trends in sorafenib and combination groups. We also noted upregulation of the LXR target genes *FASN* and *SREBF1*^24, 25^ in the GW3965 and combination groups, as expected (Figure 5C).

### Combination treatment inhibits the cell cycle and increases apoptosis

Next, we performed pathway analysis of the 915 differentially expressed genes in the combination group and found 472 gene ontology (GO) pathways to be significantly altered. The top ten altered pathways, based on gene ratio, are displayed in Figure 5D. The most dramatically altered GO pathway was regulation of cell death, with several others also relating to cell cycle and apoptosis, boxed in red in Figure 5D. When investigating steady state mRNA levels of cell cycle and apoptotic regulators, we found downregulation of many proliferation markers such as Origin Recognition Complex Subunit 1 (*ORC1*), Cell division cycle 25 A, (*CDC25A*), various cyclins (*CCNB1*, *CCNE1*), Minichromosome maintenance complex proteins (*MCM2*, *MCM5*), proliferating cell nuclear antigen (*PCNA*), cyclin-dependent kinase 2 (*CDK2*), and marker of proliferation Ki67 (*MIK67*) (Figure 6A). Moreover, we noted upregulation of many genes associated with apoptosis including Bcl-2-binding component 3 (*BBC3*), Phorbol-12-Myristate-13-Acetate-Induced Protein 1 (*PMAIP1*), Jun Proto-Oncogene, AP-1 Transcription Factor Subunit (*JUN*), TNF Receptor Superfamily Member 10b (*TNFRSF10B*), BCL2 Antagonist/Killer 1 (*BAK1*), and DNA Damage Inducible Transcript 3 (*DDIT3*) (Figure 6B). Sorafenib monotherapy also led to a moderate reduction in cell cycle and upregulation in apoptosis markers; although these effects were much more pronounced in combination therapy, again highlighting the cooperative effects of GW3965 and sorafenib.

**Figure 6:**
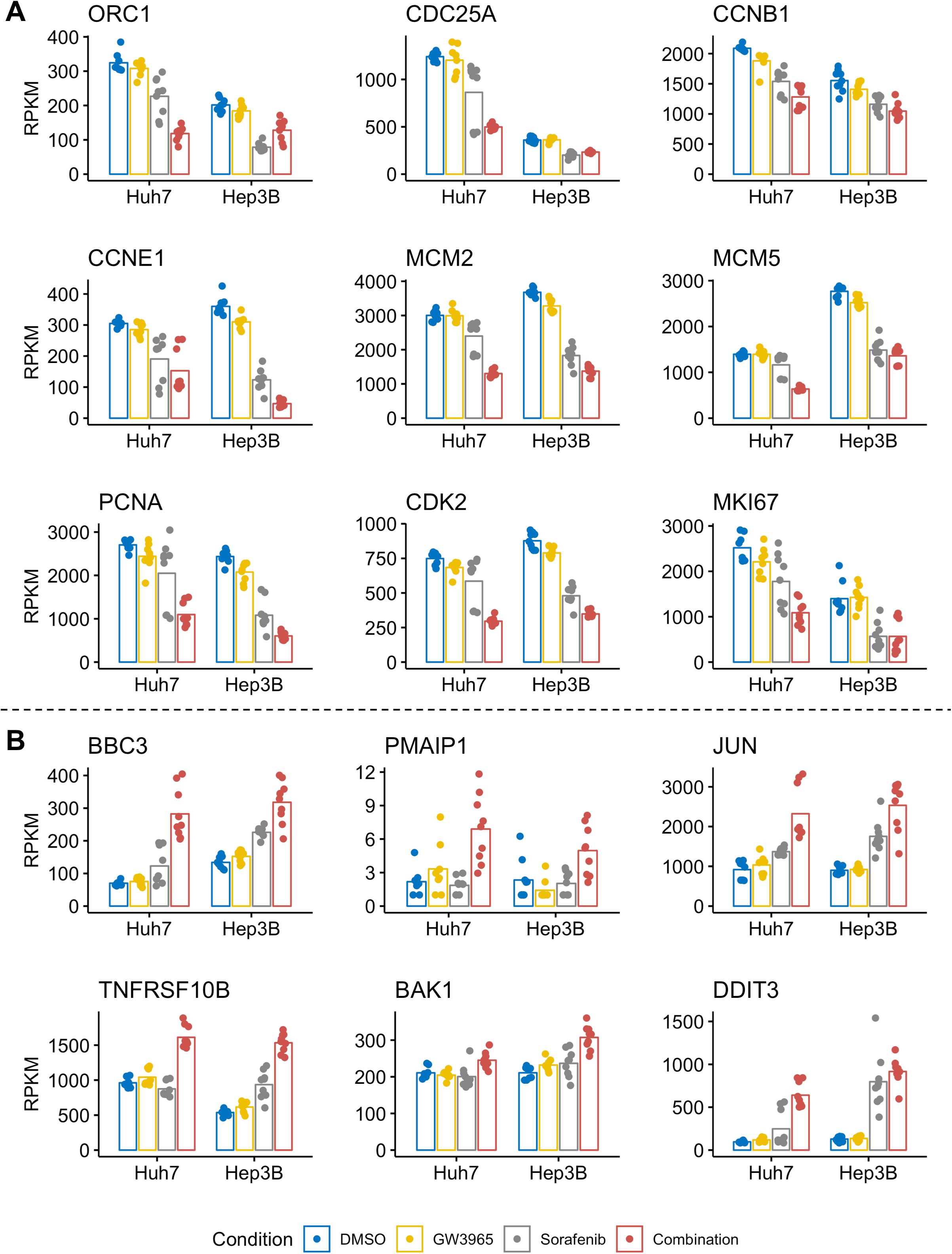
Synergistic effect of GW3965 and sorafenib in reducing cell cycle gene expression and inducing apoptosis regulators *in vitro*. (A) Reads Per Kilobase of transcript, per Million mapped reads (RPKM) of cell cycle regulators Origin recognition complex subunit 1 (*ORC1*), Cell Division Cycle 25A (*CDC25A*), Cyclin B1 (*CCNB1*), Cyclin E1 (*CCNE1*), Minichromosome Maintenance Complex Component 2 (*MCM2*), Minichromosome Maintenance Complex Component 5 (*MCM5*), Proliferating Cell Nuclear Antigen (*PCNA*), Cyclin Dependent Kinase 2 (*CDK2*), and Marker of Proliferation Ki67 (*MKI67*). (B) RPKM for apoptosis regulators BCL2 Binding Component 3 (*BBC3*), Phorbol-12-Myristate-13-Acetate-Induced Protein 1 (*PMAIP1*), Jun Proto-Oncogene (*JUN*), TNF Receptor Superfamily Member 10b (*TNFRSF10B*), BCL2 Antagonist/Killer 1 (*BAK1*), and DNA Damage Inducible Transcript 3 (*DDIT3*). The data are from Huh7 and Hep3B treated for 24 hours with DMSO (blue), GW3965 (yellow), sorafenib (gray), or combination (orange).

### Combination therapy effectively targets a patient-derived HCC

We obtained a core biopsy-derived cell line of an HCC lesion from a patient, who died 3 weeks after initiating sorafenib. We found that sorafenib monotherapy had low activity at killing this cell line, reducing viability by only 20% at the highest dose, indicating sorafenib resistance. However, when we applied the GW3965 and sorafenib combination therapy, we killed the tumor cells with similar efficiency as seen with the HCC cell lines (Figure 7A, B). Next, we extracted RNA from cells 24 hours after treatment, and observed increased transcript levels of *FASN* in GW3965 and combination treatments, and increased *GADD45B* in sorafenib and combination treatments, indicating that the combination therapy had cooperative effects on the same pathways as we observed in commercial cell lines (Figure 7C). Importantly, we found markedly reduced expression of the cell cycle promoters *CDK2, CCNE1,* and *PCNA* in the combination group (Figure 7D). Our results demonstrate that LXR agonist/sorafenib combination therapy effectively kills primary HCC cells, highlights the preclinical relevance of our studies, and suggests that drug therapy assays could be used in future personalized medicine treatment plans.

**Figure 7:**
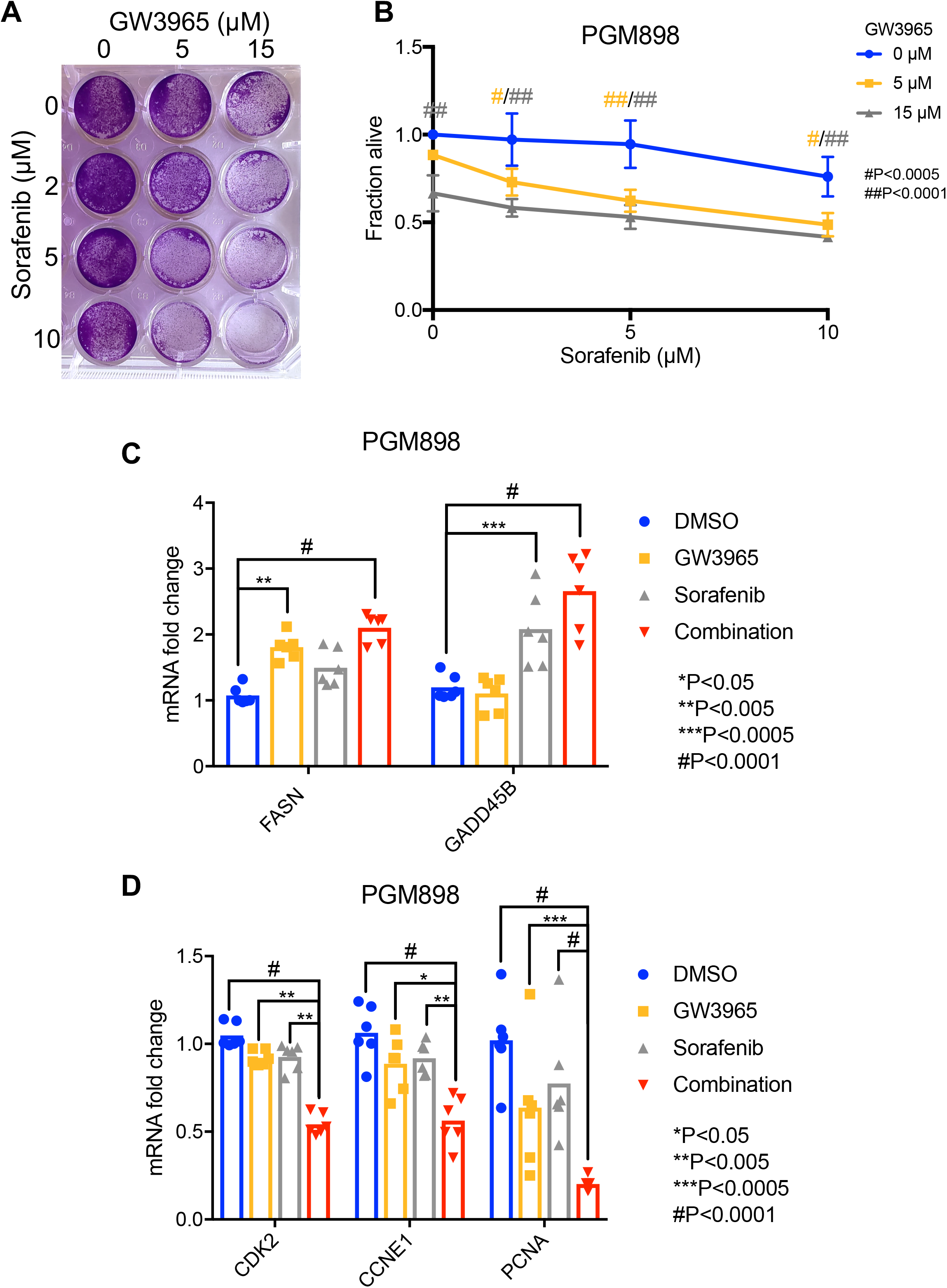
Combination treatment effectively targets patient-derived HCC *in vitro*. (A and B) Representative image and quantification (n=4) of crystal violet staining 48 hours after drug treatment. (C) RT-qPCR of FASN and GADD45B in PGM898 cells 24 hours after drug treatment (n=6). (D) RT-qPCR of CDK2, CCNE1, and PCNA in PGM898 cells 24 hours after drug treatment. Statistical analyses performed with two-way ANOVA with Tukey’s multiple comparison.

## DISCUSSION

In this study, we employed a novel genetic screen that identified LXR as a druggable target to increase the anti-tumor effect of sorafenib. Our screen utilized a previously described library of cDNAs of interest, together with the *Fah* cDNA, hydrodynamically-injected into the tail vein of *Fah*^−/−^ mice^19^. This model develops HCC within 6 weeks. We identified LXR to be disallowed in sorafenib-resistant tumors, and found that the combination of LXR agonists with sorafenib effectively kills HCC cells *in vitro* through downregulation of the cell cycle and upregulation of markers of apoptosis.

The LXR agonists GW3965 and T0901317 are known to induce conformational changes to LXR that enhances binding to coactivators. While these compounds also have an affinity for LXRß, the K_d_ values are dramatically lower for LXRα^26^. GW3965 appears to be more potent *in vitro* in combination with sorafenib, as a significant effect was observed at 48 hours (as opposed to 72 hours with T0901317) and at lower doses. Consistently across cell lines, we witnessed a significant reduction in cell viability with 5 μM GW3965 and 2 μM sorafenib. At these doses, cell viability dropped to approximately 50% in all transformed cells, but remained at 80% in the mouse hepatic cell line AML12. These findings suggest that the combination treatment preferentially kills cancer cells and spares non-cancerous cells, and could be used to reduce sorafenib dose to reduce toxicity *in vivo*.

The synergistic impact of GW3965 and sorafenib combinatorial treatment on gene expression is visualized in our heatmap in Figure 5B. While we noted cell line-specific variability, the combination treatment clearly led to similar transcriptomic changes. It is interesting that in Hep3B cells, sorafenib monotherapy led to similar changes in gene expression as the drug combination, whereas in Huh7 the sorafenib condition mostly clustered separately from the combination group. Multiple studies have found that Hep3B are more sensitive to sorafenib than Huh7^27–29^. These data, combined with our data from a sorafenib monotherapy-resistant primary HCC cell line derived from a patient, suggest that addition of GW3965 to sorafenib overcomes sorafenib resistance in HCC.

LXR regulates cholesterol and fatty acid synthesis and bile acid metabolism. A potential concern with using LXR agonists is increased fatty acid synthesis and hepatic steatosis upon LXR activation^30^. However, HCC patients with increased LXR expression have better prognosis than those with lower expression^31^, suggesting that activating LXR with an agonist is a plausible treatment option. LXR expression has been found to be upregulated in various adenocarcinomas, as compared to squamous cell carcinomas from the same tissues of origin^32^. Furthermore, LXR is expressed in macrophages and can influence macrophage-specific lipid composition^33^, begging the question of how the immune system may be impacted by LXR agonism as a cancer therapy. In various solid cancers including colon, prostate, and breast, LXR agonists have been shown to effectively target cancer through inhibition of the cell cycle^34^. Indeed, our GO pathway analysis found cell death and cell proliferation to be significantly modulated. We confirmed cell cycle downregulation through decreased expression of several cell cycle regulators, confirming that GW3965 and sorafenib impact several downstream pathways to regulate cell proliferation. Whether an increase in apoptotic regulators is a result of decreased proliferation or occurs through independent pathways remains unclear.

A wealth of pre-clinical studies, including studies in non-human primates, has shown that LXR agonists promote reverse cholesterol transport to treat atherosclerosis. This prompted two separate phase I clinical trials with LXR agonist drugs LXR-623 and BMS-852927, which were administered to more than 100 volunteers^35, 36^. Both beneficial and adverse effects on lipid metabolism occurred. For the purpose of treating cancer, the lipid changes and the side effect profile may be acceptable. In fact, there is an active clinical trial currently enrolling patients with lymphoma and a number of solid tumors (not including liver cancers) to receive an LXR agonist called RGX-104^37^.

In summary, we found that LXR agonists can enhance the activity of sorafenib in killing HCC. This result stems from our *in vivo* data demonstrating that *Nr1h3* cDNA (encoding LXR) is incompatible with the growth of MYC-driven tumors in the presence of sorafenib, and fits well with a recent study noting a correlation between LXR expression and improved HCC survival^31^. Furthermore, another publication compared the expression patterns of sorafenib-sensitive and resistant human HCC cell lines,^38^ and their data showed a significantly decreased level of LXRα expression in sorafenib-resistant cells (p < 0.01). Future studies could examine safety and efficacy of the drug combination in pre-clinical models and could perform assays with the drug combination in patient-derived HCC cells lines to personalize treatment for patients.

---

**Supplemental Figure 1: Combination therapy with LXR agonism and sorafenib.** (A and B) Quantification and representative imaging of hepatocyte-derived cell line AML12 treated with DMSO, 2 μM GW3965, 5 μM sorafenib, or GW3965 and sorafenib for 48 hours (n=10). Statistics performed using one-way ANOVA with Tukey’s multiple comparisons. (C and D) Quantification and representative images of HepG2, Hep3B, and Huh7 cells treated with varying concentrations of LXR agonist T0901317 and sorafenib for 72 hours (n=8). Statistics performed using two-way ANOVA with Tukey’s multiple comparisons.

