## Supplementary figures and images for "In vivo screen identifies LXR agonism potentiates sorafenib killing of hepatocellular carcinoma"

### Supplemental Figure 1

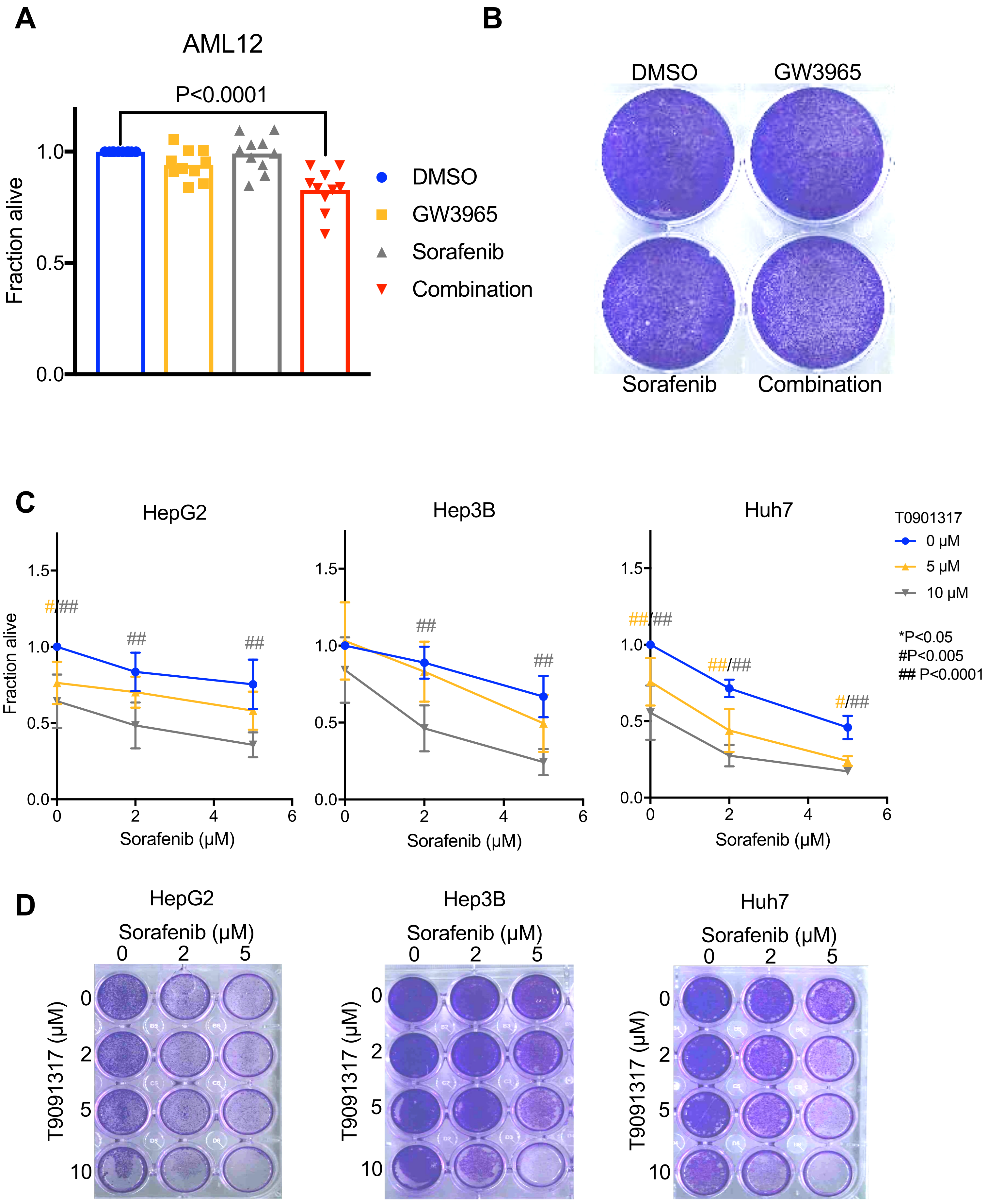
